# Bacterial contact induces polar plug disintegration to mediate whipworm egg hatching

**DOI:** 10.1101/2023.03.13.532458

**Authors:** Amicha Robertson, Joseph Sall, Mericien Venzon, Janet J. Olivas, Xuhui Zheng, Michael Cammer, Noelle Antao, Rafaela Saes Thur, Jeffrey Bethony, Peter Nejsum, Victor J. Torres, Feng-Xia Liang, Ken Cadwell

## Abstract

The bacterial microbiota promotes the life cycle of the intestine-dwelling whipworm *Trichuris* by mediating hatching of parasite eggs ingested by the mammalian host. Despite the enormous disease burden associated with *Trichuris* colonization, the mechanisms underlying this transkingdom interaction have been obscure. Here, we used a multiscale microscopy approach to define the structural events associated with bacteria-mediated hatching of eggs for the murine model parasite *Trichuris muris*. Through the combination of scanning electron microscopy (SEM) and serial block face SEM (SBFSEM), we visualized the outer surface morphology of the shell and generated 3D structures of the egg and larva during the hatching process. These images revealed that exposure to hatching-inducing bacteria catalyzed asymmetric degradation of the polar plugs prior to exit by the larva. Although unrelated bacteria induced similar loss of electron density and dissolution of the structural integrity of the plugs, egg hatching was most efficient in the presence of bacteria that bound poles with high density such as *Staphylococcus aureus*. Consistent with the ability of taxonomically distant bacteria to induce hatching, additional results suggest chitinase released from larva within the eggs degrade the plugs from the inside instead of enzymes produced by bacteria in the external environment. These findings define at ultrastructure resolution the evolutionary adaptation of a parasite for the microbe-rich environment of the mammalian gut.

## Introduction

Soil transmitted helminths (STH) are parasitic worms that affect nearly 1.5 billion people worldwide (1). The developmental maturation of certain STHs occurs in the gastrointestinal (GI) tract where the parasites encounter trillions of bacteria that are part of the gut microbiota, as exemplified by the whipworm *Trichuris*. Infection with *Trichuris* species initiates when embryonated eggs in the environment are ingested, which then hatch in the cecum and large intestine where the bacterial community is most diverse and dense (2–4). In this environment, the hatched larvae then embed themselves in the intestinal epithelium where they remain anchored as they undergo several molts to become sexually reproductive adult worms (3). Heavy worm burden in individuals infected by the human parasite *Trichuris trichiura* is associated with colitis, anaemia, and dysentery (5–8). Single doses of the available anthelmintic drugs for *T. trichiura* infections display poor efficacy with an average cure rate of less than 50% (9). A better understanding of how *Trichuris* species have adapted to their host may reveal vulnerabilities that can be targeted for intervention.

Recent studies using the murine parasite *Trichuris muris* have revealed a remarkable degree of co-adaptations with the mammalian host and the gut microbiota. Bacteria, which trigger egg hatching *in vitro*, are necessary for *T. muris* to establish infection (10–12). In turn, *T. muris* colonization alters the microbiota composition of mice (13), which we and others have shown is consequential for the host. In a model of inflammatory bowel disease (IBD), we found the type 2 immune response to *T. muris* colonization protects against intestinal inflammation by increasing Clostridiales and reducing Bacteroidales species within the gut microbiota (14). Another murine intestinal parasite *Heligmosomoides bakeri* also participates in three-way interactions with the host and microbiota including inducing an outgrowth of Clostridiales that attenuates allergic airway inflammation (15, 16). Colonization of indigenous people in Malaysia with *T. trichiura* is associated with a similar expansion of Clostridiales and reduction in Bacteroidales in the microbiota, indicating that these relationships are conserved in humans (14). Further, we found that Clostridiales species induce superior egg hatching of *T. muris* and *T. trichiura* compared with Bacteroidales species (17). These observations suggest that *Trichuris* colonization alters the microbiota composition in a manner that may be mutually beneficial for the parasite and mammalian host in certain contexts.

Given this dependence on the microbiota, egg hatching may be a point of vulnerability in the parasite lifecycle. *Trichuris* eggs are ovoid shaped with a multi-layered shell. A membrane-like outer vitelline layer covers the entire surface of the egg, and underneath is a middle layer consisting of mainly chitin and a lower lipid layer (18, 19). These eggs have two openings, one at each end, that are blocked by chitinous polar plugs (4). Although these plugs are also multilayered, they have a higher proportion of chitin in its middle layer than the rest of the shell (4). During the hatching process (eclosion), the oral spear in the anterior end of the larva pierces the outer vitelline membrane and exits through a polar plug on one side of the egg (20, 21). The role of bacteria in this process remains obscure. For the Gram-negative bacterial species *Escherichia coli*, type 1 fimbriae mediate binding to the polar ends of eggs and are necessary for optimal hatching (10). However, Gram-positive species that do not have fimbriae such as *Staphylococcus aureus* can also induce efficient hatching *in vitro* (10, 11, 17). It is unclear whether the structural events associated with hatching in the presence of Gram-negative and -positive species are similar.

In this study, we show that although many Gram-positive bacterial species are strong hatching inducers, *Enterococci* fail to trigger hatching. We also show that the ability of *S. aureus* to form high density clusters is associated with superior hatching rates and that binding of this bacterium to the egg is essential. High resolution 3D volume electron microscopy imaging revealed the ultrastructural organization of the eggshell and larva, and identified striking asymmetrical morphological changes to the polar plug regions that occur upon exposure to *E. coli* and *S. aureus*, representative Gram-negative and -positive bacteria that induce hatching. We further showed that the degree of bacterial binding to eggs is directly proportional to the hatching rate yet hatching required bacteria to be metabolically active. Finally, we show that eggs from both *T. muris* and the human pathogen *T. trichiura* harbor chitinase activity and propose that this activity plays a role in hatching by degrading the chitinous polar plug of the egg from within.

## Results

### Bacteria display species-dependent effects on *T. muris* egg hatching

To establish conditions for investigating structural changes that occur during *T. muris* egg hatching, we first investigated the degree to which bacterial taxa differ in their ability to induce hatching. Although previous studies have shown that both the Gram-negative species *E. coli* and the Gram-positive species *S. aureus* can induce hatching (10, 11), only a limited number of bacterial taxa have been examined under the same conditions and it was unclear whether the time course differs between bacterial species. We used previously optimized conditions in which bacteria cultures grown to their maximal density are added to embryonated *T. muris* eggs in a culture well and monitored over time for hatching by light microscopy under aerobic conditions (12) (Fig 1A, S1 Video). We found that *E. coli* and *S. aureus* induced comparable levels of *T. muris* egg hatching by four hours post-incubation, while untreated eggs remained unhatched (Fig 1B). However, a higher proportion of eggs were hatched in the presence of *S. aureus* at earlier time points compared with *E. coli* under these conditions (Figs 1B and E). *Pseudomonas aeruginosa* and *Salmonella enterica* Typhimurium – Gram-negative Proteobacteria related to *E. coli* – were previously shown to induce *T. muris* egg hatching under similar conditions (10, 17). To increase the number of different Gram-positive bacterial species investigated in this assay, we examined *Staphylococcus epidermidis* as a member of the same genus as *S. aureus*, and *Bacillus subtilis* and *Enterococcus faecalis* as unrelated taxa. *S. epidermidis* induced hatching at a similar rate as *S. aureus*, while *B. subtilis* also induced efficient hatching, albeit at a modestly slower rate. In contrast, hatching failed to occur in the presence of *E. faecalis*. To determine whether this was specific to *E. faecalis*, we tested another member of the genus, *Enterococcus faecium*, and found that it also did not induce hatching (Fig 1C). These differences in hatching rates cannot be explained by the culture media used to grow each bacterial species because *S. aureus* grown in each of the three media types (BHI, TSB, and LB) induced similar levels of hatching across time points (Fig 1D).

**Fig 1.**
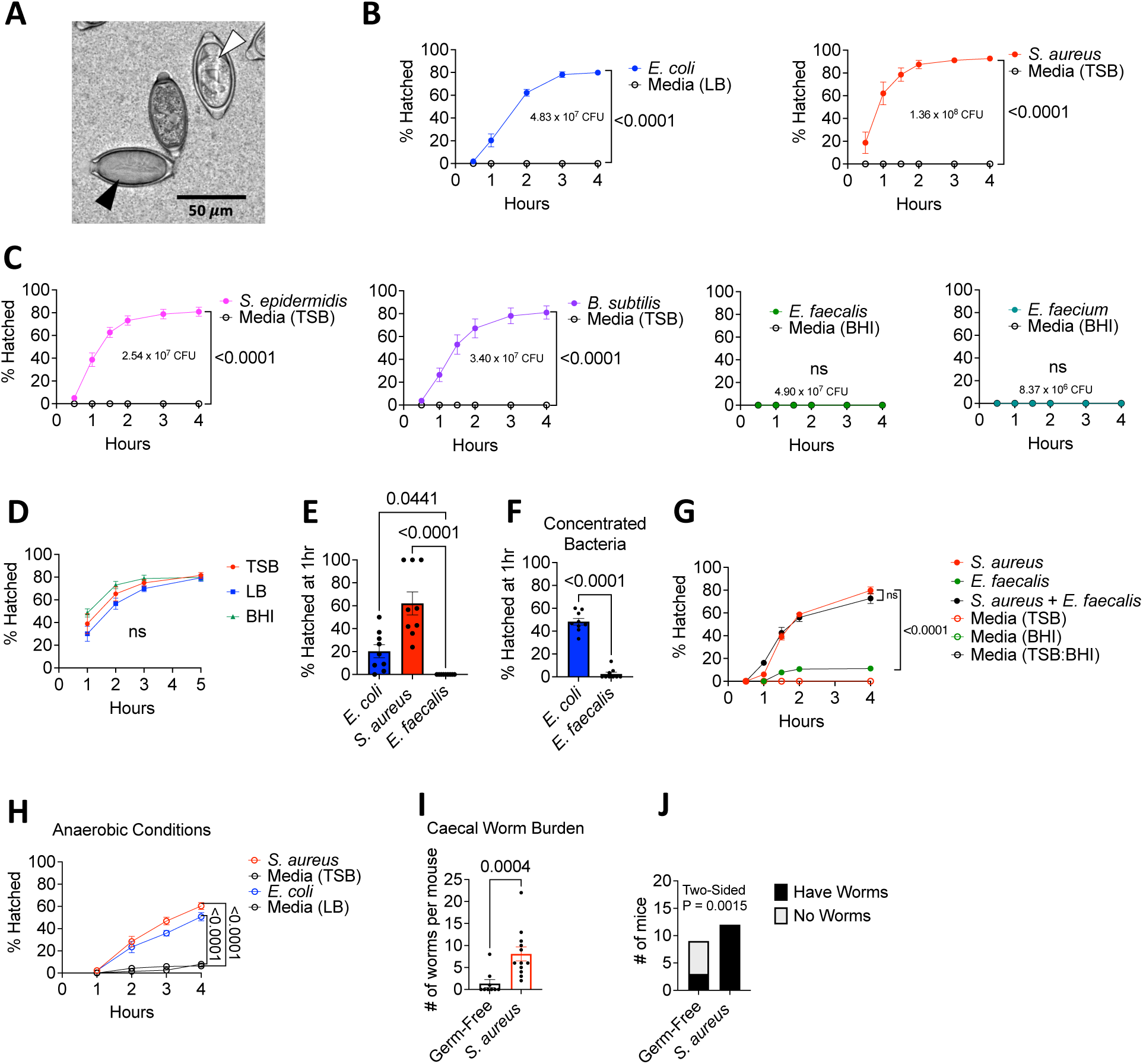
Bacteria display species-dependent effects on *T. muris* egg hatching. **(A)** Representative light microscopy image of *T. muris* eggs induced to hatch after incubation with *S. aureus* at 37℃ for 45 mins. White arrowhead denotes unhatched egg and black arrowhead denotes hatched egg. **(B)** Percent of *T. muris* eggs hatched after incubation in aerobic conditions with overnight cultures of *E. coli* and *S. aureus*, compared with their respective broth controls determined by light microscopy at indicated time points. Colony forming units (CFUs) of each bacterial species added to the eggs are indicated on the graphs. **(C)** Percent of *T. muris* eggs hatched after incubation in aerobic conditions with overnight cultures of *S. epidermidis*, *B. subtilis*, *E. faecalis* and *E. faecium* compared with their respective broth controls determined by light microscopy at indicated time points. Colony forming units (CFUs) of each bacteria added to the eggs are indicated on the graphs. **(D)** Percent of *T. muris* eggs hatched after incubation with *S. aureus* grown overnight in tryptic soy broth (TSB), Luria Bertani (LB) broth and Brain Heart Infusion (BHI) broth. **(E)** Percent of *T. muris* eggs hatched at 1 hour after 37℃ incubation with a maximum number of *E. coli, S. aureus and E. faecalis*. **(F)** Percent of *T. muris* eggs hatched at 1 hour after 37℃ incubation with an equal number (∼10^8^ CFU) of *E. coli* and *E. faecalis*. **(G)** Percent of *T. muris* eggs hatched after incubation with *S. aureus* and *E. faecalis* alone or together compared with broth controls. For the *S. aureus* + *E. faecalis* condition, bacterial cultures were grown separately overnight, and then were mixed the day of the experiment. **(H)** Percent of *T. muris* eggs hatched after incubation in anaerobic conditions with overnight cultures of *E. coli* and *S. aureus* compared with their respective broth controls determined by light microscopy at indicated time points. **(I)** Number of worms harvested from the caecum per mouse (n = 9-12 mice per group). **(J)** Proportion of germ-free and *S. aureus* monocolonized mice that had harbored adult *T. muris* worms after double-dose infection.

Data points and error bars represent mean and SEM from 3 independent repeats for (B), (C), (D), (G), and (H). Dots represent a single well and bars show means and SEM from 3 independent repeats for (E) and (F). Dots represent a single mouse and bars show means and SEM from 3 independent repeats for (I). Welch’s t test was used to compare area under the curve between each condition and its respective media control for (B), (C), and (H). Ordinary one-way analysis of variance (ANOVA) test followed by a Turkey’s multiple comparisons test was used to compare AUC of hatching induced by different conditions to each other for (D) and (G). Kruskal-Wallis test followed by a Dunn’s multiple comparisons test was used for (E). Mann Whitney test was used for (F) and (I). Fisher’s exact test was used to determine whether there was a significant association between gut microbial composition of mice and the presence of adult worms in the cecum for (J).

Consistent with the ability of staphylococci to grow as tightly packed clusters, a higher number of *S. aureus* (∼10^8^ colony forming units; CFUs) were added to eggs compared with *E. coli* (∼5x10^7^ CFUs) in the above experiments in which we normalized based on maximal growth in media. Therefore, bacterial density may explain differential *T. muris* egg hatching rates. Indeed, the difference in early hatching was eliminated when we concentrated *E. coli* to normalize the number of bacteria in the inoculum to match the maximal growth of *S. aureus* (∼10^8^ CFUs) (Fig 1F). However, increasing the number of *E. faecalis* led to negligible amounts of hatching (Fig 1F). This lack of hatching was not due to the production of an inhibitory or toxic factor by this bacterium because adding *E. faecalis* did not negatively impact *S. aureus*-mediated hatching (Fig 1G). These results show that taxonomically distant Gram-positive and -negative bacteria induce hatching with similar efficiency when the number of bacteria added to the culture are normalized, although there are Gram-positive taxa such as enterococci that are unable to induce hatching.

Given that hatching occurs predominantly in the anaerobic environment of the cecum and large intestine (2), we confirmed that *E. coli* and *S. aureus* were able to induce hatching in anaerobic conditions (Fig 1H). We previously demonstrated that monocolonization of germ-free mice with *E. coli* is sufficient to support *T. muris* development following inoculation with embryonated eggs, although the number of worms recovered is less than conventional mice (12). To determine whether a Gram-positive species can support parasite colonization, we monocolonized mice with *S. aureus* and inoculated them with two doses of 100 *T. muris* eggs. We recovered adult worms from the cecum from all *S. aureus* monocolonized mice 42 days after *T. muris* inoculation, whereas most germ-free control mice did not harbor worms (Figs 1I and J). Together with our prior findings, these results show that *E. coli* and *S. aureus*, despite their taxonomic and structural dissimilarities, are individually sufficient to induce egg hatching and promote *T. muris* colonization.

### Physical contact with egg is essential for *S. aureus*-mediated hatching

*T. muris* egg hatching is impaired in the presence of *E. coli* when direct contact is inhibited (10, 11). Given the structural and growth differences between *E. coli* and *S. aureus*, we revisited contact-dependence of hatching using the above conditions. We initially tested whether *S. aureus* secretes a soluble factor that is sufficient to induce hatching. Similar to our previous finding with *E. coli* (12), the addition of filtered *S. aureus* culture supernatant to *T. muris* eggs did not result in hatching (Fig 2A). It is possible that the presence of *T. muris* eggs is required for *S. aureus* to produce a pro-hatching factor, which we would miss in a simple supernatant transfer experiment. To address this possibility, we incubated *S. aureus* with eggs, collected and filtered the supernatant from the co-culture, and then added this egg-primed supernatant to a new batch of eggs (Fig 2B). The co-culture supernatant was also unable to induce hatching of eggs (Fig 2C). To further determine whether contact is necessary, we separated eggs by placing them inside a 0.4µm transwell insert and added bacteria to the outside well. Although moderate amounts of hatching occurred when eggs were segregated from *E. coli* in this manner, no hatching was observed when eggs were placed in transwells that were incubated with *S. aureus* (Fig 2D). These findings indicate that direct contact with eggs is required for *S. aureus* to induce hatching.

**Fig 2.**
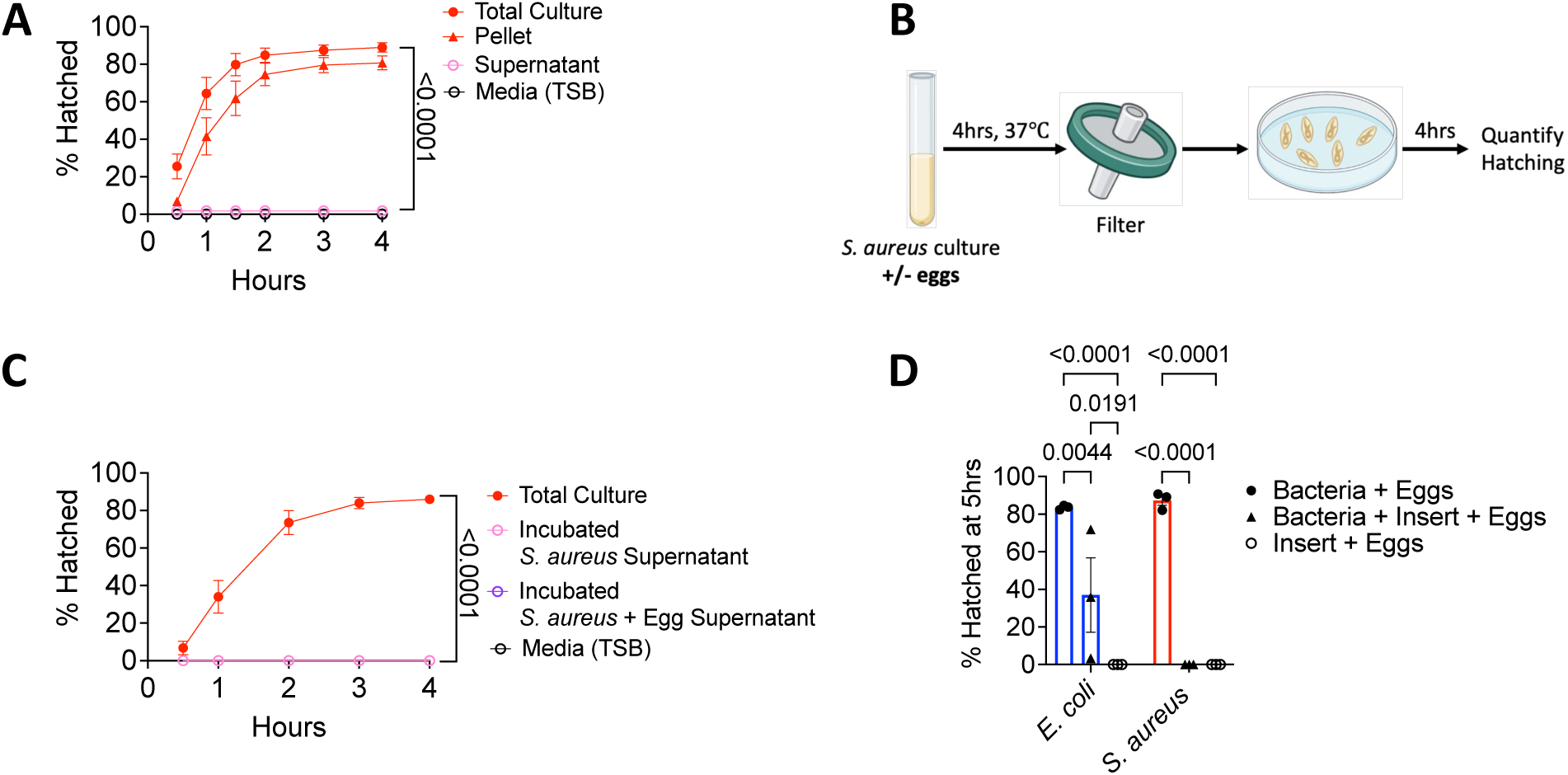
Physical contact between the bacterial cell and egg is essential for *S. aureus* mediated hatching. **(A)** Percent of *T. muris* eggs hatched after incubation with total overnight *S. aureus* culture, culture pellet resuspended in fresh media, culture supernatant filtered with a 0.22μm filter, or media control. **(B)** Experimental approach for determining whether a soluble hatching inducing factor is produced by *S. aureus* in response to exposure to eggs. Filtered supernatant from *S. aureus* grown with or without *T. muris* eggs for 4 h were transferred to a dish containing *T. muris* eggs. The ability of the supernatant to mediate egg hatching was evaluated over 4 hours. **(C)** Percent of *T. muris* eggs hatched after incubation with total overnight *S. aureus* culture or filtered supernatant obtained from *S. aureus* incubated with or without eggs as in (B) compared with media controls. **(D)** Percent of *T. muris* eggs hatched when placed in a 0.4μm transwell separated from bacteria or control media in the outer well compared with eggs incubated bacteria without transwell separation.

Data points and error bars represent the mean and SEM from 3 independent experiments for (A) and (C). Dots represent a single well and bars show means and SEM from 3 independent experiments for (D). Ordinary one-way ANOVA test followed by a Turkey’s multiple comparisons test was used to compare resuspended pellet and supernatant conditions with total O/N culture for (A) and (C). Two-way ANOVA followed by a Turkey’s multiple comparisons test was used for (D).

### Collapse of the polar plug precedes hatching mediated by bacteria

Bacteria-induced *T. muris* egg hatching has not been examined with ultrastructural resolution. We examined eggs exposed to *S. aureus* and *E. coli* by scanning electron microscopy (SEM) and were able to capture eggs at different stages in the hatching process as evidenced by larvae in mid-ejection and morphological changes to the plug not observed in untreated eggs and eggs exposed to *E. faecalis*, a poor hatching-inducing species (Figs 3A-E). For both untreated and *E. faecalis*-treated eggs, the plugs appeared as crater-like structures approximately 5 μm in diameter with an inner surface displaying a slightly rounded wrinkled morphology (Figs 3C and D). Diplococci characteristic of Enterococci were present at the plugs of eggs incubated with *E. faecalis* (Fig 3C). Untreated eggs, despite lack of hatching, had a high concentration of bacterial cells on the plugs (Fig 3D). Because eggs were harvested from adult worms isolated from the cecum of mice, these bacteria were likely derived from the mouse gut microbiota (13).

**Fig 3.**
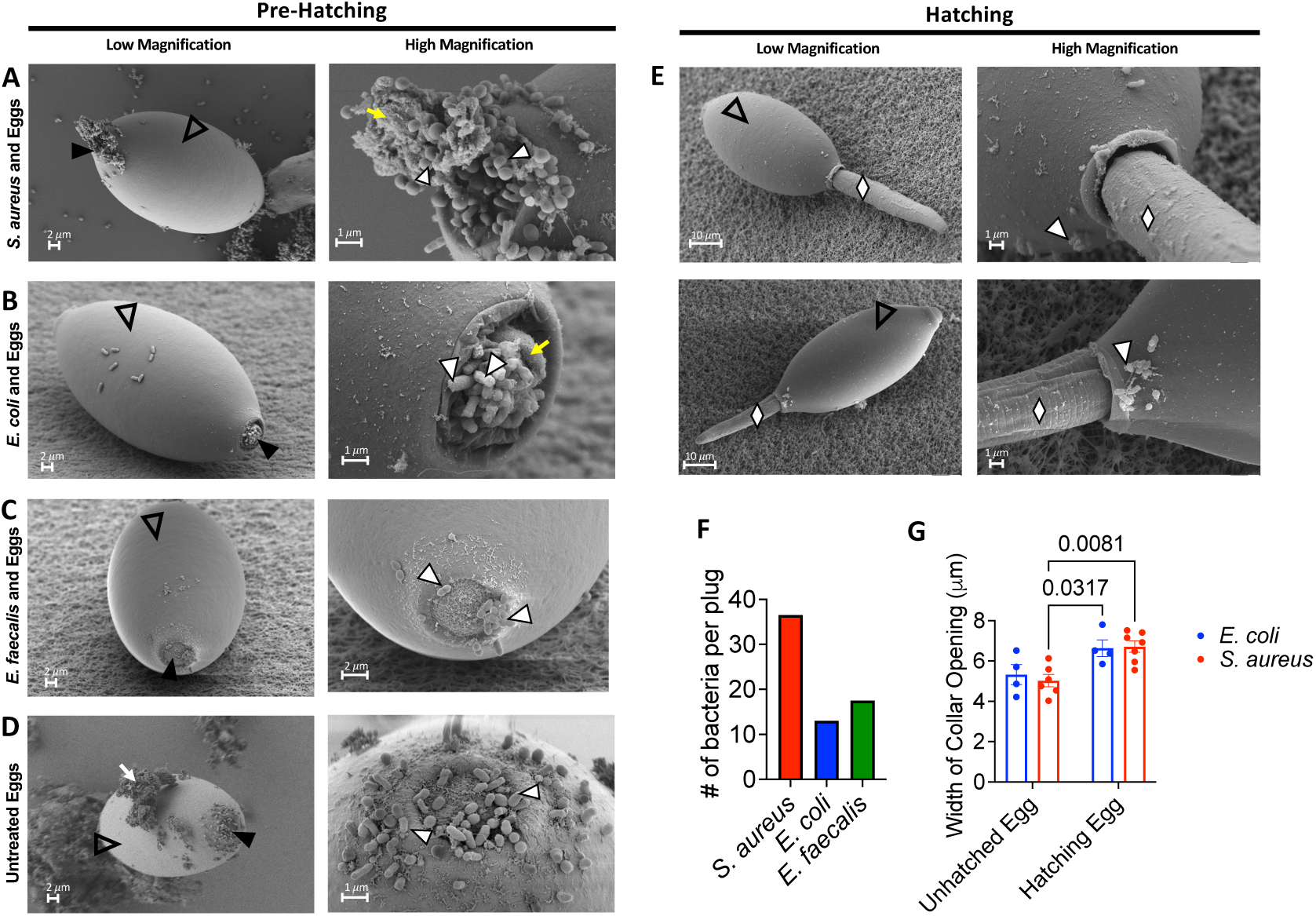
Collapse of the polar plug precedes hatching mediated by bacteria. **(A, B, C, D)** Representative low (left) and high magnification (right) SEM images of eggs (clear arrowhead) that were exposed to *S. aureus* for 1 hour (A), *E. coli* for 1.5 hours (B) and *E. faecalis* for 1 hour (C) or untreated (D). White arrowheads correspond to bacteria on polar plug regions of the eggs denoted by black arrowheads. Yellow arrow in right panel of (A) and (B) indicates woolly substance present among bacteria. Debris on untreated eggs is denoted by the white arrow (D). **(E)** Representative low (left) and high magnification (right) SEM images of hatching eggs (clear arrowhead) that were exposed to *S. aureus* for 1 hour (top) and *E. coli* for 1.5 hours (bottom). White arrowheads correspond to bacteria on polar plug regions of the eggs. Emerging larvae are denoted by white diamonds. **(F)** Number of bacterial cells visible on polar plugs of eggs incubated with *E. coli*, *S. aureus,* or *E. faecalis*. Bars showing mean from 2 eggs per condition. **(G)** Width of collar openings on eggs that were treated with either *E. coli* or *S. aureus* and were either unhatched or in the process of hatching.

For low magnification images, scale bar represents 2μm for unhatched egg in (A), (B), (C) and (D) and 10μm for hatched egg in (A) and (B). For high magnification images, scale bar represents 1μm for (A), (B) and (D) and 2μm for (C). Dots represent a single plug and bars show mean and SEM of collar sizes from 4-7 eggs per condition for (G). Two-way ANOVA followed by a Turkey’s multiple comparisons test was used for (G).

Eggs incubated with *S. aureus* and *E. coli* were enriched for the presence of bacteria at their plugs (Figs 3A and B), which displayed a collapsed morphology distinct from untreated eggs (Fig 3D) and eggs incubated with *E. faecalis* (Fig 3C). *S. aureus* and *E. coli* were densely clustered within or near depressions. Additionally, there was a wooly substance that was closely associated with the bacteria on the plug region (Figs 3A and B, right). The vitelline membrane of the plug was also visible and resembled a deflated balloon (Fig 3B, right). Bacteria, while present sporadically across the eggshell, were more densely packed on the plug region than other regions. Examination of eggs incubated with *S. aureus* and *E. coli* in which worms were captured mid-ejection showed that, in both instances, larvae exited the egg through the plug, rupturing it along with the outer vitelline layer in the process (Fig 3E). In these eggs in which larvae were partially out of the shell, bacteria were found near the polar plug region (Fig 3E, top and bottom).

We observed more individual bacterial cells per disrupted plug when eggs were incubated with *S. aureus* than other conditions, likely reflecting their capacity for growing in clusters. However, the number of visible *E. faecalis* cells on the polar plug region was comparable to the number of *E. coli* cells on the structurally distinct polar plug, suggesting that differences in the density of bacterial species at the plug are not sufficient to explain why Enterococci are poor hatching inducers (Fig 3F). Additionally, the diameter of the collar region on eggs in the process of hatching was generally wider than on eggs exposed to bacteria, but that had not yet begun to hatch, consistent with the previously described observation that plugs swell during the hatching process (21) (Fig 3G). These results show that exposure of eggs to strong hatching-inducing bacteria is associated with a collapse and loss of structural integrity of the polar plugs.

### Hatching-inducing bacteria trigger disintegration of the polar plug

Advances in the volume electron microscopy technique serial block face-scanning electron microscopy (SBFSEM) present an opportunity to gain additional insight into the above structural changes that we observed in eggs exposed to *S. aureus* and *E. coli.* Preparation of samples for ultrastructural electron microscopy imaging normally involves chemical fixation, staining, dehydration and then finally embedding of the specimen in resin (22). However, the impermeability of the eggshell made preservation of the egg contents using routine chemical fixation and high pressure freezing and freeze substitution methods difficult (23). To overcome this challenge, we used a microwave assisted sample preparation method to increase the penetration of fixative and stains into the egg, which enabled identification of structures corresponding to the eggshell, polar plugs, and the larvae (Figs 4A-C, S2 Video). The eggshell thickness was comparable between both bacteria-treated eggs, 2.17*μ*m and 2.18µm for *E. coli* and *S. aureus*-exposed eggs, respectively (Figs 4A-C). Cells and other structures within the larvae and the cuticle confining the larvae were visible. Characteristic granules of the larval intestinal tract were observed, and we propose that those granules contain lipids based on the low electron density of the contents (Fig 4A, S2 Video) (20). The plug on one end (top) displayed a more granular and less electron dense morphology compared with the contralateral plug (bottom) for both *S. aureus*- and *E. coli*-exposed eggs. The anterior end of the larva was closer to the top plug that was granular and pointed towards the bottom plug in both cases (Figs 4A-C, S2 Video). 3D renderings generated from the data collected showed that both *S. aureus* and *E. coli* were sporadically present over the entire surface of the egg and enriched at the plugs (Figs 4B and C respectively, right; S2 Video). In the case of *S. aureus* exposed eggs, a large aggregate of a characteristic grape-like cluster of cocci was visible on one of the poles, versus the shell where most bacteria were present as either single cells or in pairs (Figs 4B, far right).

**Fig 4.**
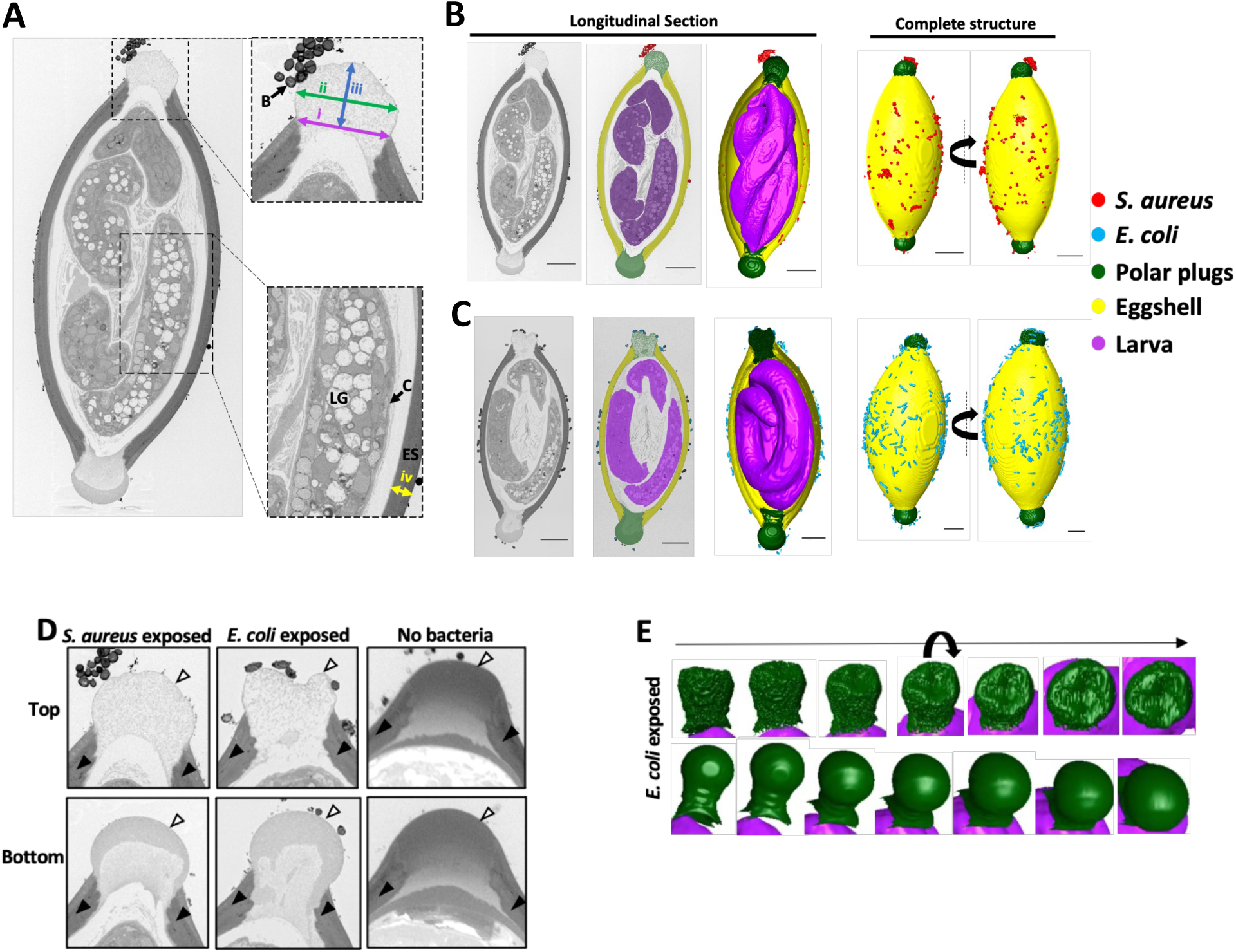
Bacteria induce asymmetric disintegration of the polar plug. **(A)** Representative electron micrograph of a longitudinal cross-section of a *T. muris* egg exposed to *S. aureus* for 1 hour. Image shows granules containing lipids (LG) within the larva, eggshell (ES) and larval cuticle (C). Distances measured are also indicated. The width of the collar opening (i), width of the widest part of the plug (ii), the height of the plug (distance from the top of the collar to the top of the plug) (iii) and the eggshell thickness (iv) were measured for all conditions. Insets show the regions boxed in black dotted line. **(B, C)** Representative electron micrographs and 3D reconstructions from SBFSEM data of a *T. muris* egg exposed to *S. aureus* for 1 hour (B) and *E. coli* for 1.5 hours (C). For the longitudinal section, the original micrograph (left), the micrograph with color overlays indicating segmented egg components and bacteria (middle) and 3D reconstruction of the egg (right) are shown. For the complete structure, two different angles are shown (left and right). *S. aureus* (red), *E. coli* (blue), polar plugs (green), eggshell (yellow) and larvae (purple) are all shown. Scale bars represent 10*μ*m. **(D)** High magnification images of polar plugs from (B) and (C) on eggs exposed to *S. aureus* and *E. coli* compared with equivalent regions from an egg untreated with bacteria. Outer vitelline layer is denoted by white arrowheads and eggshell is denoted by black arrowheads. **(E)** 3D reconstruction of polar plugs on eggs exposed to *E. coli*. Multiple angles are shown (left to right).

For *Trichuris* eggs not treated with bacteria, we found that the fixative did not penetrate through the intact shell even with the microwave assisted fixation method, precluding structural analysis of the larva. However, we were able to obtain high resolution images of the eggshell and pole regions of untreated eggs for comparison with bacteria-treated eggs. Eggshells were 2.31µm thick, similar to bacteria-treated eggs (Figs 4A-D). Consistent with previous reports analyzing eggs in non-hatching conditions (18, 21), we found that the polar plugs appeared as uniformly electron dense structures rounded towards the outer surface for egg samples that were not exposed to hatching-inducing bacteria (Fig 4D, right). While being rounded, the height of the plug of the untreated egg (Fig 4A) did not extend far beyond the top of the collar on this untreated egg (3.29µm top, 3.49µm bottom. These intact plugs also contained an electron dense outer vitelline layer that was continuous with the rest of the shell and overlayed a similarly electron dense chitinous layer enclosed by an electron dense, ridged collar (Fig 4D, right).

In contrast, eggs incubated with *S. aureus* or *E. coli* displayed plugs that were morphologically distinct from untreated egg plugs. First, they were much less electron dense and non-uniform in electron density. As noted above, these changes were asymmetric. For both bacteria, one plug completely lost structural integrity and appeared more granular and disintegrated than the contralateral plug on the same egg (Fig 4D). Instead of a smooth rounded surface, these plugs displayed indentations on the surface, which was more pronounced for the *E. coli*-treated egg in which the plug had a large depression (Figs 4D and E; S2 Video). The contralateral pole was more intact compared with the top pole but was less electron dense compared with plugs on the egg not exposed to bacteria. There were two distinct regions of electron density for these contralateral poles exposed to *S. aureus* or *E. coli*, with an outer region of moderate density encompassing an inner less electron dense area (bottom) (Fig 4D, left and middle). In the *S. aureus* condition, the plug with the disintegrated morphology corresponded with the side associated with the bacterial clusters. Although different in electron density, both plugs on eggs incubated with bacteria were swollen and extended beyond the width of the collar region. Specifically, plugs on *E. coli* exposed eggs were 9.51µm and 10.43µm in width while their respective collars were 8.87µm and 8.68µm in diameter for the top and bottom, respectively (Fig 4D). Similarly, plugs on *S. aureus* exposed eggs were 9.46µm and 10.39µm wide while their respective collars were 8.62µm and 9.46µm wide (Fig 4D). Moreover, the heights of the plugs were larger on eggs exposed to *E. coli* and *S. aureus* (4.47µm top and 7.40µm bottom, and 5.63µm top and 6.51µm bottom, respectively) than on the untreated egg (Fig 4D).

We repeated the SBFSEM experiment with new bacteria-exposed egg samples to determine whether we can capture additional stages of plug disintegration. In this second run of SBFSEM on *S. aureus-*exposed eggs, one of the plugs (pole 1, S1 Fig A) displayed similar morphology as the contralateral (bottom) plug on the *S. aureus-*exposed egg in Fig 4D, while the other plug (pole 2, S1Fig A) had a similar electron density to the outer region of the pole 1 plug but was uniformly electron dense. Additionally, the second run of SBFSEM on *E. coli-*exposed eggs showed plugs with a uniform and high electron density that were not swollen, similar to the plugs seen on eggs that were not treated with bacteria (S1 Fig B). Lastly, in the second run on *S. aureus* exposed eggs, we observed that the larva appeared to be directly interacting with the inner surface of pole 1 (S1 Fig C). This result could be representative of additional and earlier stages of plug degradation.

Together, these results indicate that exposure to hatching inducing bacteria is associated with substantial morphological changes of the polar plugs, including swelling and a decrease in structural integrity.

### Degree of bacterial binding is proportional to the rate of *T. muris* egg hatching

The strict requirement of physical binding for *S. aureus-*induced hatching and the striking images showing that this bacterial species was densely bound to the plug motivated us to quantitatively define the nature of this interaction. Towards this end, we used spinning-disk confocal microscopy to quantify the location and number of bacteria by incubating eggs with a *S. aureus* strain that expressed GFP (Fig 5A). At 30 minutes post-incubation, we found that the pole region contained a substantially higher proportion of GFP+ structures compared with the sides of the eggs (Fig 5B). Our cross-species comparisons (Fig 1) suggest that hatching efficiency is dependent on the number of bacteria incubated with the eggs. Consistent with this observation, when we added serial dilutions of *S. aureus* to eggs, we observed that the rate of hatching was proportional to the concentration of bacteria (Fig 5C). To test whether this relationship between bacterial concentration and hatching rates reflected binding events, we quantified the number and location of bacteria associated with eggs incubated with dilutions of *S. aureus-*GFP. The percentage of eggs associated with GFP+ puncta and the number of GFP+ puncta per egg after a 30-minute incubation were proportional to the concentration of *S. aureus* added to the culture (Figs 5D-F). In contrast to the higher concentration conditions in which almost all GFP+ structures were associated with poles, the proportion of *S. aureus* bound to the poles versus other parts of the eggshell was reduced at lower concentrations (Figure 5G). These results are consistent with the ultrastructural analyses, suggesting that the degree of *S. aureus* binding to the polar plug region determines the rate of hatching.

**Fig 5.**
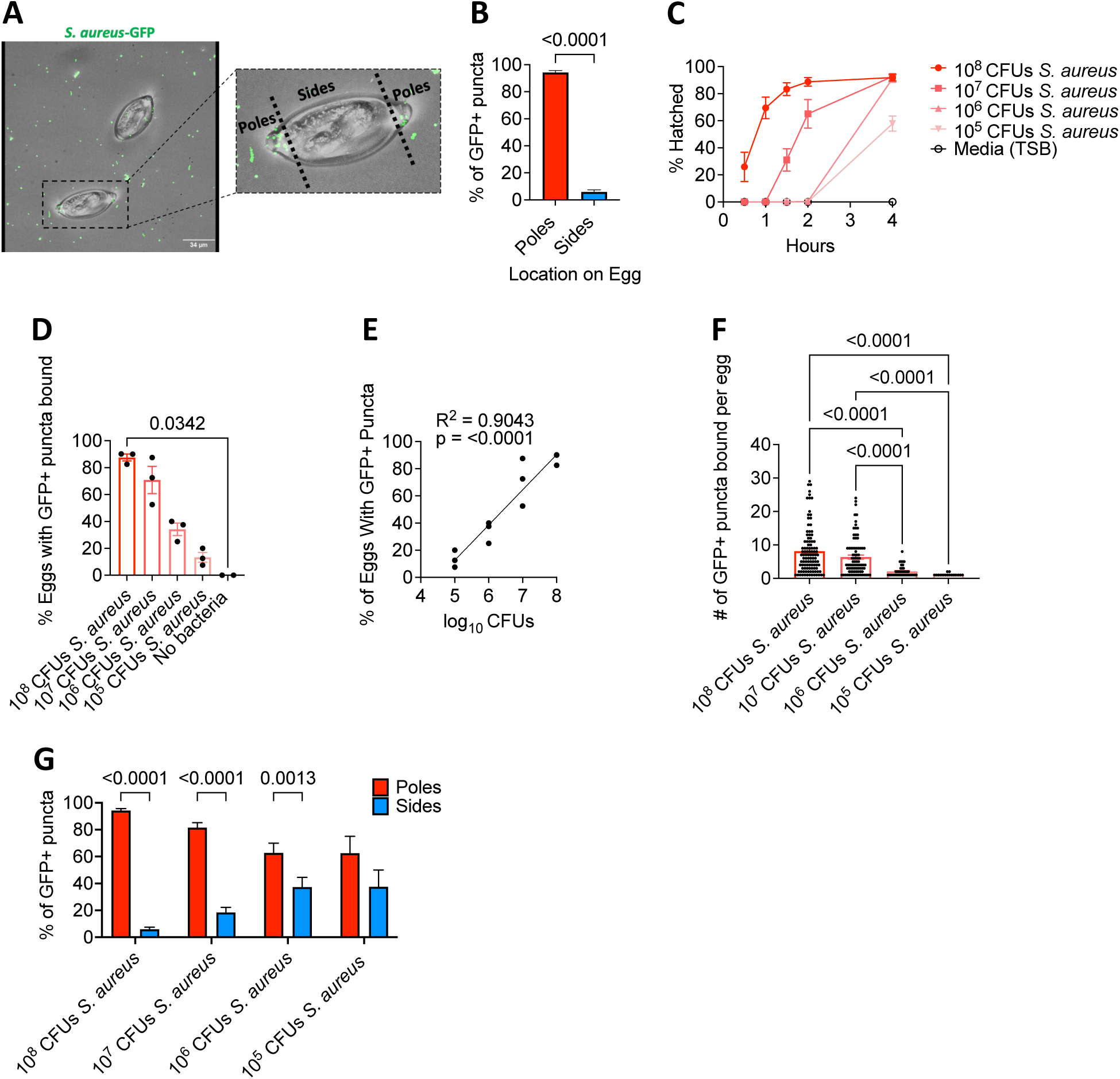
Degree of bacterial binding is directly proportional to the rate of *T. muris* egg hatching. **(A)** Representative confocal microscopy image of eggs incubated with *S. aureus* GFP for 30 minutes. Regions defined as poles and sides are indicated. Scale bar represents 34μm. **(B)** Percent of GFP+ puncta present on the poles versus the sides of the eggs. **(C)** Percent of *T. muris* eggs hatched after incubation with 10-fold dilutions of overnight *S. aureus* culture ranging from approximately 10^5^ – 10^8^ CFU (n = 3). **(D)** Percent of eggs associated with GFP+ puncta after incubation with 10-fold dilutions of overnight *S. aureus-*GFP culture ranging from approximately 10^5^ – 10^8^ CFU. **(E)** Correlation analysis comparing log_10_ CFUs of bacteria used and percent of eggs with GFP+ puncta present. **(F)** Number of GFP+ puncta bound per egg from (D). **(G)** Percent of GFP+ puncta present on the poles of the eggs versus the sides of the eggs from (D).

Bars and error bars show means and SEM from 3 independent repeats for (B), (D), (F) and (G). Data points and error bars represent mean and SEM from 3 independent repeats for (C). Dots represent percentage of 40 eggs that were GFP+ from a single experiment for (D) and (E) and number of GFP+ puncta found on a single egg for (F). Mann-Whitney test was used for (B). Kruskal-Wallis test followed by a Dunn’s multiple comparisons test was used for (D) and (F). Simple linear regression was performed for (E). Two-way ANOVA followed by a Sidak’s multiple comparisons test was used for (G).

### Bacterial metabolic activity is essential for *S. aureus-*mediated hatching

Egg hatching was low or completely absent within the first 30 minutes post-incubation with *S. aureus* (Figs 5C and 6A). Given our results showing a direct relationship between bacterial binding and hatching rate, it is possible this delay in hatching reflects the time required for bacteria to bind eggs. However, even at 0 minutes post-incubation with *S. aureus*-GFP, the majority of eggs were bound by bacteria based on their association with GFP+ puncta (Figure 6B). By 10 minutes post-incubation, the proportion of eggs bound to bacteria and the number of bacterial cells bound to each egg reached the maximum value (Figs 6B and C). *S. aureus* was consistently enriched at plugs, and the ratio of bacteria bound to plugs versus other regions of the shell was similar across time points (Fig 6D). These findings indicate that bacterial binding is not the rate limiting step for hatching induction.

**Fig 6.**
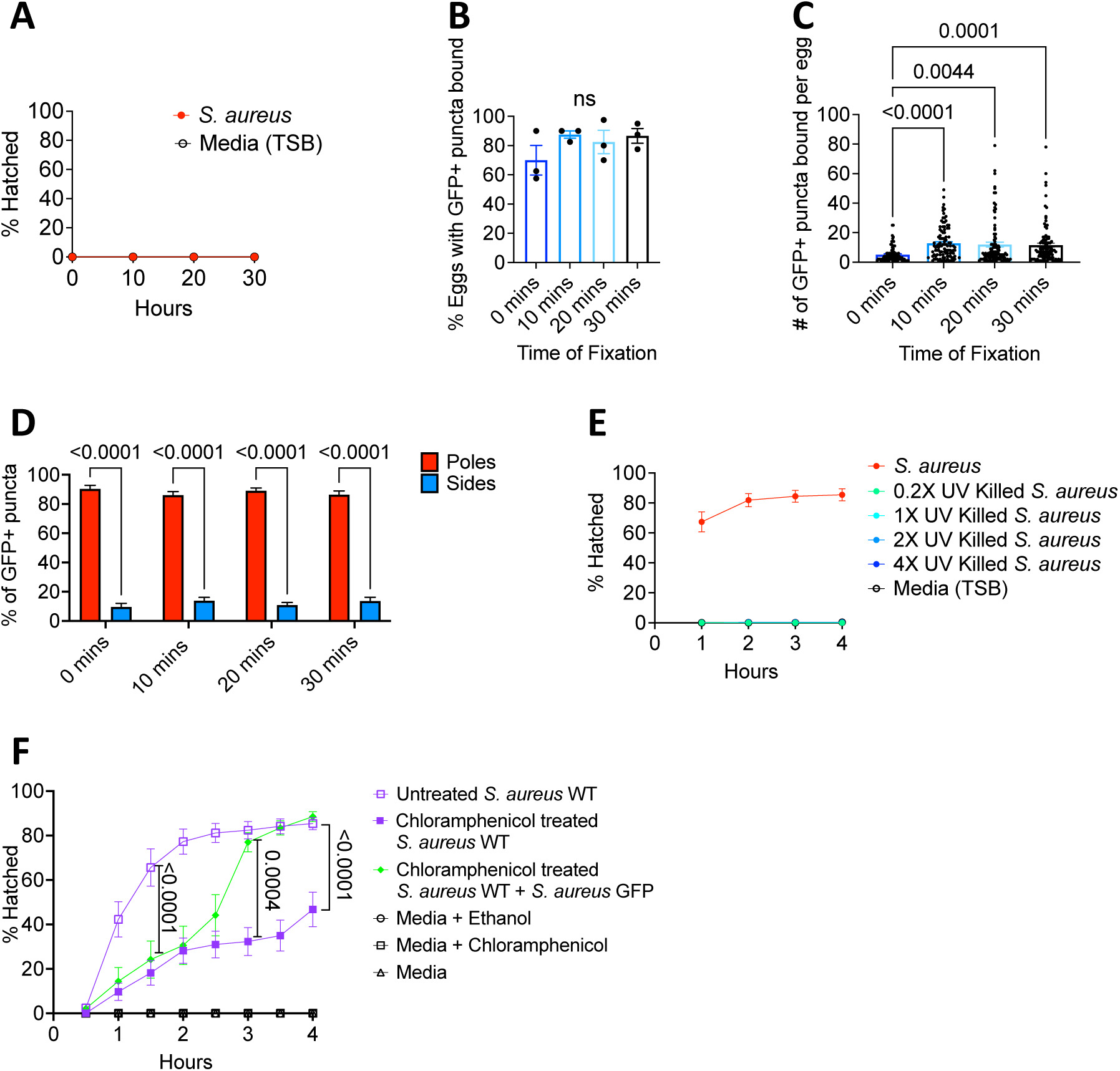
Bacterial metabolic activity is essential for *S. aureus* mediated hatching. **(A)** Percent of *T. muris* eggs hatched after incubation with 10^8^ CFU overnight *S. aureus* culture at time points indicated. **(B)** Percent of eggs that had GFP+ puncta bound after incubation with 10^8^ CFU of overnight *S. aureus* culture for different lengths of time. **(C)** Number of GFP+ puncta bound per egg after incubation with 10^8^ CFU of overnight *S. aureus* culture for different lengths of time. **(D)** Percent of GFP+ puncta present on the poles of the eggs versus the sides of the eggs after incubation with 10^8^ CFU of overnight *S. aureus* culture for different lengths of time. **(E)** Percent of *T. muris* eggs hatched after incubation with UV killed overnight *S. aureus* culture. **(F)** Percent of *T. muris* eggs hatched after incubation with untreated *S. aureus*, chloramphenicol (CAM)-treated *S. aureus* (100μg/ml), and CAM-treated *S. aureus* together with CAM-resistant *S. aureus* GFP for 4hrs. *S. aureus* GFP was spiked-in at the 2hr time point.

Bars and error bars show means and SEM from 3 independent repeats for (B), (C), and (D). Data points and error bars represent mean and SEM from 3 independent repeats for (A), (E) and (F). Dots represent mean percentage of 40 eggs that were GFP+ from a single experiment for (B) and number of GFP+ puncta found on a single egg for (C). Kruskal-Wallis test followed by a Dunn’s multiple comparisons test was used for (B) and (C). Two-way ANOVA followed by a Sidak’s multiple comparisons test was used for (D). Ordinary one-way ANOVA followed by Turkey’s multiple comparisons test was used to compare AUC of different conditions tested for (F).

We previously showed that hatching required metabolically active *E. coli*, specifically arginine biosynthesis (12). Consistent with a requirement for viability, UV-killed *S. aureus* cells failed to induce egg hatching (Fig 6E). Also, the bacteriostatic antibiotic chloramphenicol (CAM), which inhibits protein translation in *S. aureus* (24, 25), reduced bacteria-mediated hatching (Fig 6F). This impaired hatching was not due to an effect of CAM on the egg itself because spiking-in *S. aureus-*GFP that is CAM-resistant at 2 hours after the beginning of the assay immediately restored hatching in the presence of CAM (Fig 6F). In conclusion, egg binding and metabolic activity are both required for *S. aureus* to mediate hatching.

### *Trichuris* eggs harbor chitinase activity

The biochemical process involved in *Trichuris* egg hatching is unknown. Hatching of eggs from the STH *Ascaris lumbricoides* is mediated by a parasite-derived chitinase enzyme (26). However, *A. lumbricoides* hatching occurs independently of bacteria. Thus, we explored the possibility that bacteria provided the chitinase for *T. muris* eggs to hatch. *S. aureus*, for example, produces degradative enzymes that have chitinase activity (27). To test this hypothesis, we used a *S. aureus* mutant with a deletion in a gene encoding a chitinase-related protein, which we refer to as CRP. Hatching occurred at a similar rate in the presence of *S. aureus Δcrp* and wild-type, indicating that CRP is not required (Fig 7A). In fact, S*. aureus* mutants deficient in all the major secreted proteases (*S. aureus ΔaurΔsspABΔscpAspl::erm*) (28) as well as a *S. aureus* mutant lacking the three major lipases (*S. aureus ΔgehAΔgehBΔgehE*) (29) retained the ability to induce hatching with similar efficiency as wild-type bacteria (Figs 7B and C). These findings raise the possibility that the enzymatic activity responsible for the disintegration of the poles is derived from the parasite rather than bacteria.

**Fig 7.**
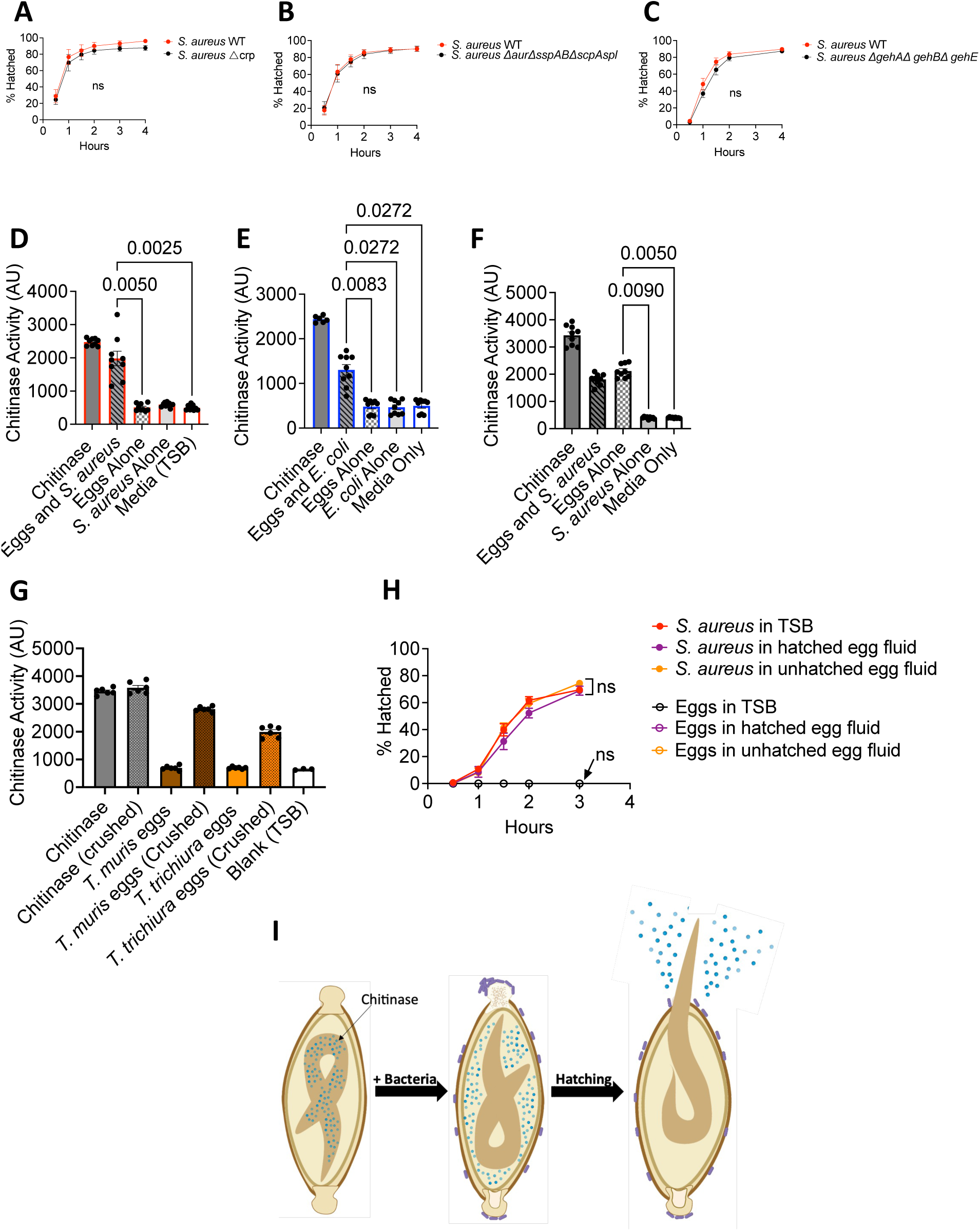
*Trichuris* eggs harbor chitinase activity. **(A-C)** Percent of *T. muris* eggs hatched after incubation with overnight *S. aureus* WT, *S. aureus Δ*crp (A), *S. aureus ΔaurΔsspABΔscpAspl::erm* (B) and *S. aureus ΔgehAΔgehBΔgehE* (C) culture. **(D, E)** Amount of fluorescence detected in wells containing eggs hatched in response to *S. aureus* (D) and *E. coli* (E). AU = arbitrary unit. 5mU of stock chitinase were used as a positive control. **(F)** Amount of fluorescence detected in wells containing crushed eggs that were exposed to *S. aureus* for 20 minutes. 0.02mU of stock chitinase were used as a positive control. **(G)** Amount of fluorescence detected in wells containing crushed *T. muris* and *T. trichiura* eggs. 0.02mU of stock chitinase were used as a positive control. **(H)** Percent of *T. muris* eggs hatched after incubation with fluid from hatched eggs (purple open circles) and bacteria resuspended in fluid from hatched eggs (purple filled circles). Fluid from unhatched eggs was used as a control (yellow filled and open circles). **(I)** Graphical depiction of proposed mechanism.

Data points and error bars represent means and SEM from 3 independent repeats for (A), (B), (C) and (H). Dots represent fluorescence detected in a single well and bars and error bars show means and SEM from 2-3 independent repeats for (D), (E), (F) and (G). Welch’s t test was used to compare area under the curve between mutant *S. aureus* strains and WT *S. aureus* for (A), (B) and (C). Kruskal-Wallis test followed by a Dunn’s multiple comparisons test was used for (D), (E), (F) and (G). Ordinary one-way ANOVA followed by Turkey’s multiple comparisons test was used to compare AUC of different conditions tested for (H).

Chitinase is released into the media during *A. lumbricoides* egg hatching (26). If *T. muris* also uses chitinase to degrade the polar plug from the inside, we should be able to detect chitinase activity in the post-hatching fluid, similar to *A. lumbricoides*. Using a previously described assay in which a substrate produces a fluorescent product when cleaved by chitinase (30), we detected chitinase activity in media from eggs incubated with *S. aureus*, but not in media containing untreated eggs or *S. aureus* alone (Fig 7D). Release of chitinase was not specific to *S. aureus*-mediated hatching as chitinase activity was also detected in media from eggs incubated with *E. coli* (Fig 7E). To determine whether chitinase within *T. muris* eggs is induced by exposure to bacteria, we measured chitinase activity in crushed eggs with or without pre-incubation with *S. aureus*. Chitinase activity was detected in the media containing crushed eggs that were exposed to bacteria as well as in eggs that were not exposed to bacteria (Fig 7F). Thus, eggs harbor chitinase activity independent of bacteria treatment. This experiment also rules out the possibility that bacteria were the source of the chitinase activity. We detected chitinase from crushed *T. trichiura* eggs (Fig 7G), indicating that the human parasite also produces this enzyme.

Lastly, we tested whether media containing chitinase released from *T. muris* eggs induce hatching of intact eggs from the external environment (“outside-in”). Fluid from hatched eggs was obtained by incubating eggs with bacteria for 4hrs and then filtering the supernatant. Fluid obtained from unhatched eggs was included as a negative control. We observed that fluid from hatched eggs did not induce hatching and was also unable to enhance *S. aureus*-mediated hatching (Fig 7H). These results suggest that enzymes likely mediate hatching from the inside rather than from the outside (Fig 7I).

## Discussion

In this study we illuminated structural events that occur during bacteria-mediated hatching of *T. muris* eggs by using multiscale microscopy techniques. Notably, we overcame technical challenges to apply SBFSEM to generate high resolution 3D images of eggs in the process of hatching. This technique along with SEM showed that, despite being taxonomically distant species with significant structural and biological differences, *E. coli* and *S. aureus* induce similar morphological changes in the polar plugs of eggs. These structural changes include swelling of the plugs, and disintegration of the plug material, resulting in loss of structural integrity of the plugs that appear as dips and depressions on the plug surface.

A previous study describing the egg hatching cascade showed that prior to hatching, the head of the larva inserts into the plug space and then uses its stylet to tear through the vitelline membrane of the plug and exit the egg (21). Therefore, it is possible that bacteria-induced plug disintegration plays an important role in hatching by either hollowing out the space in the collar region to accommodate the larval head or by reducing the density of the plug to facilitate movement into the plug space. The difference in electron density we observed between contralateral plugs on the same egg might be indicative of plugs at different stages of degradation, with the less dense plugs being further along in the disintegration process. This model is supported by our comparison of plugs from different eggs, which highlighted the heterogenous degrees of collapse and granularity. The fact that a loss of electron density is observed during hatching induction by two different bacterial species increases our confidence that dissolution of the integrity of the polar plugs is an essential step during hatching. A mechanism in which disintegration of the poles is rate-limiting and occurs in an asynchronous manner would explain why bacteria-induced egg hatching is also asynchronous.

The strict requirement for physical binding between *S. aureus* and *T. muris* eggs, and the general absence of information regarding how Gram-positive bacteria mediate hatching, motivated us to further investigate hatching of eggs exposed to this bacterium by confocal microscopy. Although the number of bacterial cells bound to the poles was a critical determinant of hatching efficiency, physical association was not sufficient. Also, although we observed *E. faecium* bound to egg poles, the two *Enterococcus* species we tested failed to induce significant amounts of hatching. Similar to our previous findings that bacterial byproducts were necessary for *E. coli*-mediated hatching (12), *S. aureus* metabolic activity contributed to hatching. These results support a requirement for both bacterial binding and bacterial products.

Given that the polar plug predominantly consists of chitin, we initially considered the possibility that binding led to local production of chitinase derived from bacteria, thereby degrading the pole from the outside. However, hatching remained efficient in the presence of *S. aureus* mutants lacking candidate enzymes including a homolog of chitinase. Additionally, fluid collected from hatched eggs was unable to induce hatching of unexposed eggs, suggesting that polar plug disintegration is occurring via an “inside-out” mechanism where the larva itself produces the enzymes that facilitate eclosion (Fig 7I). Consistent with this possibility, we detected chitinase within the eggs. A mechanism by which the larva within the egg is the producer of degradative enzymes would explain why unrelated bacterial taxa can induce hatching – it would be unlikely that all the bacterial species shown to induce hatching possess the same enzyme necessary to dissolve the chitinous plug. This model would be further supported if a conserved bacterial product were shown to induce hatching. Such a factor has not been identified. The lack of a genetic system to interrogate *T. muris* and the impermeable nature of the eggshell also pose challenges for elucidating detailed mechanism. Therefore, technical advances may be required to confirm the role for worm-derived chitinase and examine how products released from bacteria relate to this chitinase activity. Although bacteria exposure is not required for the presence of chitinase activity, it may influence the location of the enzymatic activity (Fig 7I). It is possible, for instance, that bacterial exposure triggers the release of chitinase from vesicles within the worm and into the perivitelline space in the egg, as suggested for *H. bakeri* (31).

Gram-positive species are a prominent component of the gut microbiota and certain taxa are enriched in *Trichuris*-colonized individuals (17). *S. aureus* and other staphylococci colonize the gut of humans, especially early in life as the microbiota develops (32, 33). It is possible that microbiota composition contributes to the susceptibility of young children to high worm burden and disease. Differences in microbiota composition may also partially explain why helminth infections tend to be aggregated in the host population, with a minority of the population harboring a majority of the worms (34–36). Our findings indicate that, although not all bacteria induce hatching of *T. muris* eggs, those that mediate hatching cause similar structural events to occur at the polar plug. We suggest that targeting enzymatic activity involved in plug disintegration would be difficult if it initiates from within an impermeable eggshell that evolved to be resilient to the external environment. Instead, understanding the chemical and physical interaction between bacteria and eggs may uncover a vulnerability in the life cycle of the parasite that is suitable for intervention.

## Materials and methods

### Experimental model details

#### Parasite maintenance

Stock eggs of *Trichuris muris* E strain (14) were propagated and maintained in the NOD.Cg-*Prkdc^scid^*/J (Jax) mouse strain as previously described (37). Each egg batch was confirmed to hatch at ≥ 90% *in vitro* using method below and WT *S. aureus* before use in subsequent experiments.

Stock eggs of *Trichuris trichiura* were provided by the *Trichuris trichiura* egg Production Unit (TTPU) located at the Clinical Immunology Laboratory at the George Washington University. Whipworm eggs were isolated from the feces of a chronically infected human volunteer following a qualified standard procedure that includes a modified Simulated Gastric Fluid (SGF) method. After isolation, the eggs were stored for two months in flasks containing sulfuric acid (H_2_SO_4_) maintained at 25-30°C in a monitored incubator. Once embryonation was achieved, the eggs were transferred to a locked and monitored refrigerator at 2-8°C until further use. Controls for the manufacturing process involved: (i) tests for viability (hatching), which confirmed that more than 80% of the eggs were viable; (ii) species confirmation by polymerase chain reaction (PCR); and (iii) evaluation of the microbiological burden determined by bioburden testing by an outside-certified laboratory.

### Bacterial Strains

*Escherichia coli* strain used was a kanamycin resistant transformant of strain BW25113 kindly provided by E. Jane Hubbard (NYU Grossman School of Medicine) (12). *Staphylococcus aureus* strain USA300 LAC clone AH1263 (38), *S. epidermidis* (ATCC12228) and *Enterococcus faecalis* OG1RF were provided by V. J. Torres (NYU Grossman School of Medicine) and were originally sourced from Alex Horswill (University of Colorado Anschutz Medical Campus), Eric Skaar (Vanderbilt University) and Lynn Hancock (University of Kansas) respectively. AH1263 was used as wild-type (WT) *S. aureus* unless otherwise specified. *Bacillus subtilis* was sourced from the ATCC (ATCC 6633). *E. faecium* Com15 was kindly provided by Howard Hang (Scripps Research). Frozen glycerol stocks (30% glycerol) of all bacteria were prepared. Glycerol stocks of *E. coli* was streaked onto Luria Bertani (LB) kanamycin 25 mg/ml plates (NYU Reagent Preparation Core), *S. aureus, S. epidermidis* and *B. subtilis* were streaked onto Tryptic Soy Agar (TSA) plates (NYU Reagent Preparation Core) and *Enterococcus* species were streaked onto bile esculin azide (BEA) plates (Millipore) and incubated overnight at 37°C. Single colonies of *E. coli, Staphylococcus* species and *Enterococcus* species were then spiked into 5mls of LB broth, Tryptic Soy Broth (TSB) and Bacto Brain Heart Infusion (BHI) broth (BD) respectively unless otherwise specified and grown overnight at 37°C with shaking at 225 rpm. *B. subtilis* was also spiked into TSB and was grown overnight under the same conditions. To quantify colony forming units, we performed serial dilutions of liquid culture in sterile PBS and plated on LB (Kan) agar for *E. coli*, TSA for *Staphylococcus species* and *B. subtilis* and BEA *for Enterococcus species*.

### S. aureus mutants

The *S. aureus ΔaurΔsspABΔscpAspl::erm* mutant*, S. aureus ΔgehAΔgehBΔgehE* mutant*, S. aureus Δcrp mutant* (and its parental strain LAC (ErmS)) and *S. aureus* GFP strain were all provided by V. J. Torres (NYU Grossman School of Medicine) and were originally sourced from A. Horswill (University of Iowa) (28), Francis Alonzo (University of Illinois at Chicago), Lindsey Shaw (University of South Florida), and the V. Torres lab respectively*. S. aureus* GFP strain used was a chloramphenicol resistant transformant of strain AH-LAC USA300 transformed with a pOS1 vector containing superfolder gfp with a sarA promoter (39). Frozen glycerol stocks (30% glycerol) of all bacteria were prepared. Glycerol stocks of *S. aureus ΔaurΔsspABΔscpAspl::erm, S. aureus ΔgehAΔgehBΔgehE* mutant*, S. aureus Δcrp mutant* and *S. aureus* GFP were streaked onto TSA plates with 5 μg/ml of erythromycin, 25 μg/ml Kan/ 25 μg/ml neomycin (Neo), 50 μg/ml Kan/ 50 μg/ml Neo and 10 mg/ml of chloramphenicol respectively (NYU Reagent Preparation Core and V. Torres lab). Single colonies of all species were spiked into 5mls of TSB and grown overnight at 37°C with shaking at 225 rpm. To quantify colony forming units, we performed serial dilutions of liquid culture in sterile PBS and plated on TSA plates.

### Germ-free Mice

Germ-free (GF) C57BL/6J mice were bred and maintained in flexible-film isolators at the New York University Grossman School of Medicine Gnotobiotics Animal Facility. Absence of fecal bacteria was confirmed monthly by evaluating the presence of 16S DNA in stool samples by qPCR as previously described (40). Mice were transferred into individually ventilated Tecniplast ISOcages for infections to maintain sterility under positive air pressure. Female mice 6-11 weeks of age were used for all experiments in this study. All animal studies were performed according to protocols approved by the NYU Grossman School of Medicine Institutional Animal Care and Use Committee (IACUC) and Institutional Review Board.

## Method details

### In vitro hatching of T. muris eggs

*T. muris* eggs were hatched *in vitro* by mixing 25 μL of embryonated eggs at a concentration of 1 egg/1 μL suspended in sterile water with 10 μL of overnight bacterial culture and 15 μL sterile media in individual wells of a 48 well plate. Media control wells contained an additional 10 *μ*l of sterile media instead of bacterial culture. Plates were incubated at 37°C and observed every 10 mins, 30 mins and/or hour for 4-5 hrs on the Zeiss Primovert microscope to enumerate the percentage of hatched eggs by counting hatched and embryonated unhatched eggs in each well. Unembryonated eggs, which lack visible larvae and have disordered contents, were excluded due to their inability to hatch. For anaerobic hatching specifically, plates were incubated at 37°C in an anaerobic chamber and a separate 48-well plate was used per timepoint as the plate needed to be removed from the anaerobic chamber to count hatching each timepoint. Experiments utilizing transwell inserts (Millicell) were performed as previously described (11). For experiments where cell-free supernatant was used, supernatant and cells were isolated by centrifugation and filtration through a 0.22 μm syringe filter or after wash with autoclaved PBS, respectively. Incubation with embryonated eggs was performed by adding 30 *μ*L *T. muris* eggs to 5 mL of bacterial culture and incubating at 37°C for four hours. *E. coli* and *E. faecalis* were concentrated by centrifuging 4 ml of each and then resuspending the pellets in 400 *μ*l of their respective media. For the co-incubation experiment, 2 ml of *S. aureus* overnight culture was added to 2 ml of *E. faecalis* overnight culture and then the mixture was added to eggs.

### *T. muris in vivo* infection in germ-free mice

Female germ-free C57BL/6J mice were monocolonized at 6–11 weeks of age by oral gavage with ∼1 × 10^9^ colony forming units (CFU) of *S. aureus*. A subculture of *S. aureus* was made by diluting overnight culture 1:100 in TSB and then incubating at 37°C while shaking at 225 rpm for ∼4 hrs, until 1 × 10^9^ CFU/mL was reached. 5ml of subculture was pelleted by centrifugation at 3480 rpm for 10 min and washed once with sterile 1x PBS. Pellets were then resuspended in 500 μL sterile 1x PBS and mice were inoculated by oral gavage with 1x10^9^ CFU in a volume of 100 μL. Inoculum was verified using dilution plating of aliquots.

7 and 28 days later, mice were infected by oral gavage with ∼100 embryonated *T. muris* eggs. 14 days after the second gavage of *T. muris* eggs, mice were euthanized, and individual worms were collected and enumerated from the cecal contents of mice.

### Electron microscopy

#### Scanning electron microscopy

*T. muris* eggs were incubated with *S. aureus* or *E. faecalis* for 1hr, *E. coli* for 1.5 hr or bacteria free media for 1 hr. All eggs were fixed with 2% paraformaldehyde (PFA), 2.5% glutaraldehyde in 0.1 M sodium cacodylate buffer (CB, pH 7.2) at 4°C for a week. ∼50 eggs were then loaded into 12 mm, 0.44 mm transwells (#PICM01250, Millipore Sigma) to avoid losing eggs during sample processing. The eggs were washed 3 times with PBS, post fixed with 1% osmium tetroxide (OsO_4_) in aqueous solution for 1 hour, then dehydrated in a series of ethanol solutions (30%, 50%, 70%, 85%, 95%) for 15 mins each at room temperature. Eggs were then dehydrated with 100% ethanol 3 times for 20 mins each. The eggs were critical point dried using the Tousimis Autosamdri®-931 critical point dryer (Tousimis, Rockville, MD), put on the SEM stabs covered with double sided electron conducted tape, coated with gold/palladium by the Safematic CCU-010 SEM coating system (Rave Scientific, Somerset, NJ), and imaged with the Zeiss Gemini300 FESEM (Carl Zeiss Microscopy, Oberkochen, Germany) using secondary electron detector (SE_2_) at 5 kV with working distance (WD) 18.3 mm.

#### Serial block face-scanning electron microscopy

*T. muris* eggs were incubated with *S. aureus*, *E. coli* or bacteria-free media and fixed as described above. For further sample processing, we adopted a previously described protocol (41) and made modifications based on personal communications with Rick Webb (Queensland University) using a PELCO Biowave (Ted Pella Inc., Redding, CA) for microwave processing. The detailed sample processing steps are listed in table 1. Samples were embedded with Spurr resin (Eletron Microscopy Sciences, Hatfield, PA) between ACLARE using a sandwich method.

**Table 1.**
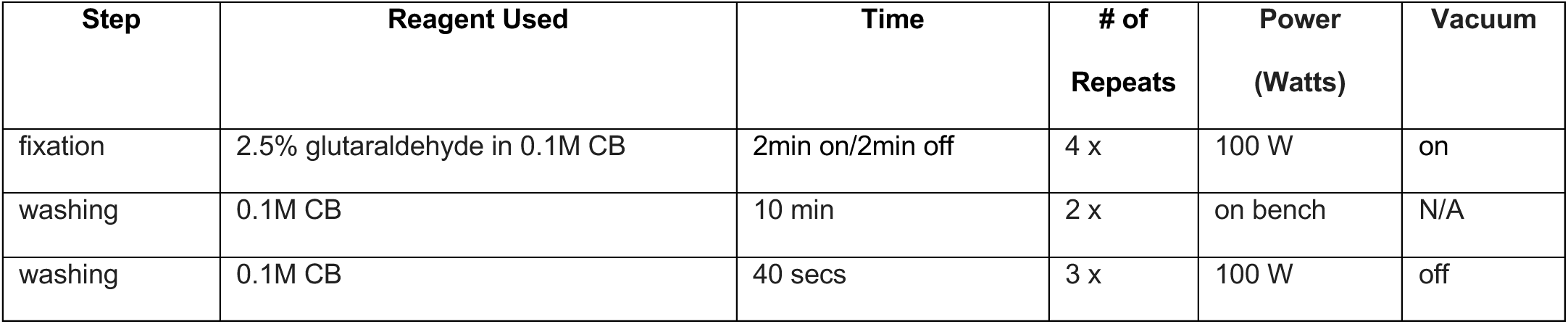
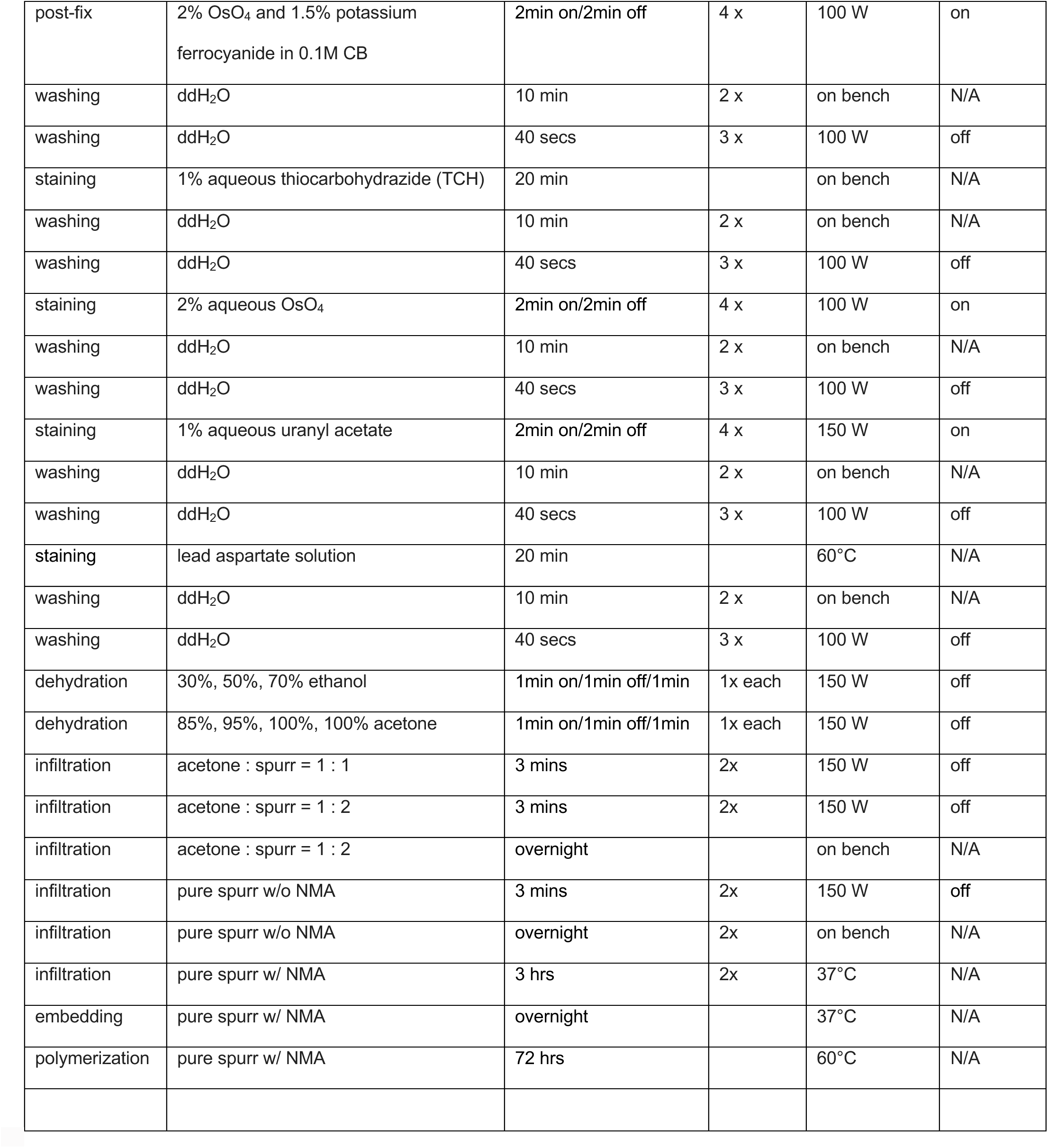
Sample preparation process for SBF-SEM imaging.

For SBF-SEM imaging, the sample block was mounted on an aluminum specimen pin (Gatan, Pleasanton, CA) using silver conductive epoxy (Ted Pella Inc.) to electrically ground the block. The specimen was trimmed again and sputter coated with 10 nm of gold (Rave Scientific, Somerset, NJ). Serial block face imaging was performed using a Gatan OnPoint BSE detector in a Zeiss Gemini 300 FESEM equipped with a Gatan 3View automatic microtome unit. The system was set to cut sections with 100 nm thickness, imaged with a pixel size of 12 nm and a dwell time of 3.0 μs/pixel, with each frame sized at 50 x 90 μm. SEM beam acceleration was set at 1.5 keV and Focus Charge Compensation gas injection set at 12% (7.9E-04mBar) to reduce charging artifacts. Images of the block face were recorded after each sectioning cycle with a working distance of 6.6 nm. Data acquisition and sectioning were automatically controlled using Gatan Digital Micrograph software to manage imaging parameters. A stack of 250 slices was aligned and assembled using ImageJ. Semi-automated segmentation and video renders were generated with ORS Dragonfly 4.1 (Object Research Systems, Montréal, QC).

### Optical microscopy

#### Live Imaging

20 μL of overnight *S. aureus*-GFP culture were added to 50 μL of embryonated eggs (1 egg/1 μL) and 30 μL sterile media in a 35 mm Petri dishes with No. 1.5 coverglass at the bottom. Mixture was then imaged using a Nikon 40x N.A. 1.3 oil immersion objective lens on a Nikon Eclipse Ti microscope. An environmental chamber set at 37°C and a lens heater set at 45°C were both used to observe hatching for approximately 45 minutes.

#### Binding Assay

An *in vitro* hatching assay was prepared as described above using *S. aureus-*GFP. After incubating the plate at 37°C, eggs with bacteria were fixed using 4% PFA, 0.5% glutaraldehyde in PBS at room temperature for 1 hr. Plate contents were then transferred to 1.5 ml Eppendorf tubes. Eggs in this mixture were pelleted by centrifugation at 1000 rpm for 5 mins with a slow brake speed (4). Samples were then washed twice with PBS and then resuspended in PBS for imaging.

Fixed samples were imaged on 35 mm Petri dishes with No. 1.5 coverglass at the bottom using a Nikon 60x N.A. 1.4 oil immersion objective lens on a Nikon Eclipse Ti microscope. Number of puncta on the sides and on the poles were enumerated by eye.

### UV killing of *S. aureus*

2 ml overnight of *S. aureus* culture were added to 8 ml of sterile PBS in a 150mm Petri dish. The dish was then placed uncovered in a UV Stratalinker 2400 (Stratagene) for 1 hr. Killed bacteria were then transferred to a 15 ml conical tube and then centrifuged at 3480 rpm and 4°C for 10 mins. Pellets were then resuspended in different amounts of sterile TSB to obtain bacterial suspensions of different concentration that were then used in an *in vitro* hatching assay as described above.

### Chloramphenicol treatment of *S. aureus*

Overnight cultures of *S. aureus* were centrifuged at 3480 rpm and 4°C for 10 mins, resuspended in fresh, sterile TSB and then incubated with 100 μg/ml of chloramphenicol or 4% ethanol (vehicle control) on ice for 45 mins as previously described (25). Treated bacteria was then used in an *in vitro* hatching assay described above. After 2 hrs of incubation, 10 μl of untreated chloramphenicol resistant *S. aureus* (*S. aureus-*GFP) was added to the wells of the hatching assay and hatching was monitored for 2 more hours.

### Chitinase detection assay

*In vitro* hatching assay was performed in a 96-well black-walled, clear, flat-bottom plate by adding 12.5 μl of eggs to 7.5 μl of sterile media and 5 μl of overnight bacterial culture and incubating the plate at 37°C for 4 hrs. 6.25 μl of 4-Methylumbelliferyl *β*-D-N,N’,N’- triacetylchitotrioside hydrate (4MeUmb, Millipore Sigma) were then added to hatching assay wells as well as an additional well containing 25 μl of chitinase from *Streptomyces griseus* (positive control, Millipore Sigma). After incubating at 37°C for 1 hr, reaction was stopped by adding 6.25 μl of 1M glycine/NaOH (NYU Reagent Preparation Core) to all wells. Plate was then read at 355/450nm ex/em by using a SpectraMax M3 plate reader (Molecular Devices) (30).

Chitinase activity within eggs exposed to bacteria was measured by first preparing an *in vitro* hatching assay in a 48-well plate as described above and incubating it at 37°C for 20 mins. 16 wells per condition were prepared. Well contents for each condition were then pooled together into 1.5 ml Safe-lock tubes (Eppendorf) with 1.0 mm diameter glass beads (BioSpec Products). Tubes were then homogenized by performing 4, 20 s homogenization cycles at 4.5 m/s using a bead beater. 25 μl of homogenate was then added to a 96-well black-walled, clear, flat-bottom plate and chitinase detection assay was performed as described above.

Chitinase activity within *Trichuris* eggs that were not exposed to bacteria was measured by first centrifuging equal numbers of eggs (500g, 3 mins, brake=3 for *T. trichiura* and 4000rpm, 5 mins, brake=4 for *T. muris*) and then resuspending them in sterile TSB. Eggs were then homogenized and chitinase detection assay performed as described above. In both homogenization experiments, chitinase from *Streptomyces griseus* (Millipore Sigma) was also homogenized as a positive control.

### Determining effect of hatching fluid on eggs

An *in vitro* hatching assay was prepared in a 48-well plate as described above and incubated at 37°C for 4 hrs. 36 wells per condition were prepared. Well contents for each condition were pooled together, centrifuged at 3480 rpm for 10 mins, and then supernatant was filtered as described above and added to new eggs in another *in vitro* hatching assay. Hatching was measured over time.

Additional *S. aureus* overnight cultures were also centrifuged at 3480 rpm for 10 mins and the remaining pellet was resuspended in the same filtered hatching assay fluid obtained above. Resuspended *S. aureus* was then added to new eggs in another *in vitro* hatching assay and hatching was measured over time.

### Quantification and statistical analysis

The number of repeats per group is annotated in corresponding figure legends. Significance for all experiments was assessed using GraphPad Prism software (GraphPad). Specific tests are annotated in corresponding figure legends. p values are also annotated on the figures themselves. A p value of ns or no symbol = not significant.

## Acknowledgements

We would like to thank Margie Alva, Juan Carrasquillo, and David Basnight for their help in the NYU Gnotobiotic Facility, and the NYU Reagent Preparation service for providing bacterial media. We thank Gira Bhabha, Damian Ekiert, Shruti Naik and members of the Cadwell and P’ng Loke Labs for their constructive comments and technical assistance. Additionally, we would like to thank David Hall, Nathan Schroeder and E. Jane Hubbard for their assistance with identifying larval structures in SBFSEM images. Figures 2B and 7I were created using BioRender.com. Lastly, we thank Chris Petzold and Jason Liang and the NYU Microscopy Laboratory for their consultation and timely preparation of the electron microscopy work. This core is partially supported by the NYU Cancer Center Support Grant NIH/NCI P30CA016087, and the Gemini300 FESEM was supported by NIH S10 OD019974.

## Supporting information

**S1 Fig.**
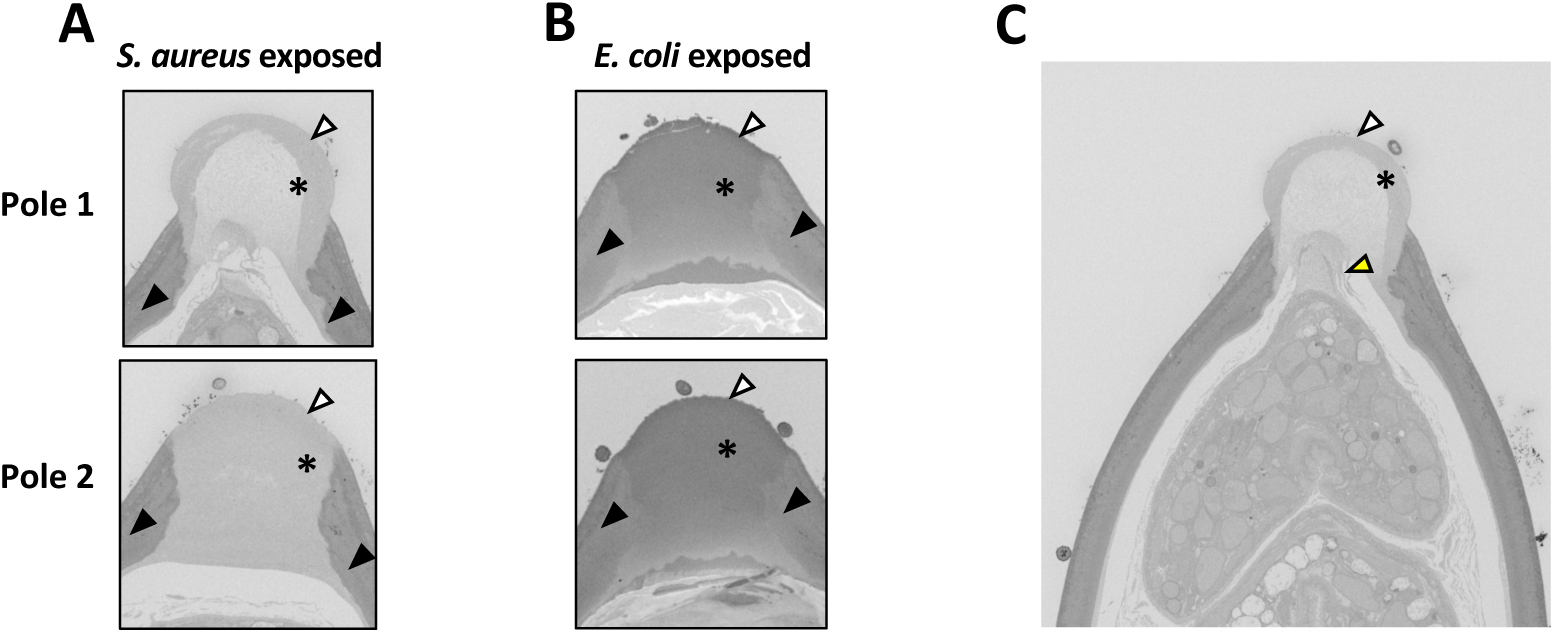
Plugs on eggs from bacteria exposed samples show different stages of disintegration, related to Fig 4. **(A, B)** Representative electron micrograph of a section of polar plugs (black asterisk) on eggs exposed to *S. aureus* (left) or *E. coli* (right). Images of Pole 1 (top) and Pole 2 (bottom) were collected from the same egg. Outer vitelline layer is denoted by white arrowheads and eggshell is denoted by black arrowheads. **(C)** Representative electron micrograph of a section of polar plug (black asterisk) on an egg exposed to *S. aureus.* Outer vitelline layer is denoted by the white arrowhead and point of contact between inner surface of the plug and the larva is denoted by the yellow arrowhead.

**S1 Video. Live imaging of *T. muris* egg hatching.** *T. muris* eggs were incubated with 2 x 10^10^ cfu/ml of *S. aureus* and imaged continuously at 37℃ for 45 mins. White arrows indicate eggs that hatch. Eggs were imaged using a 40x N.A. 1.3 objective.

**S2 Video: 3D reconstruction of a *T. muris* egg incubated with *E. coli* for 1.5 hours at 37°C from SBFSEM**

Video begins with slices through the egg and bacteria being shown. The 3D reconstruction of the larva (purple) and the polar plugs (green) is then revealed. One plug curves inward and the other plug appears rounded. Lastly, the reconstructed shell (yellow) and bacteria (blue) are shown.

